# A split methyl halide transferase AND gate that reports by synthesizing an indicator gas

**DOI:** 10.1101/2020.06.11.146191

**Authors:** Emily M. Fulk, Dongkuk Huh, Joshua T. Atkinson, Margaret Lie, Caroline A. Masiello, Jonathan J. Silberg

## Abstract

It is challenging to detect microbial reactions in highly opaque or autofluorescent environments like soils, seawater, and wastewater. To develop a simple approach for monitoring post-translational reactions within microbes situated in environmental matrices, we designed a methyl halide transferase (MHT) fragment complementation assay that reports by synthesizing an indicator gas. We show that backbone fission within regions of high sequence variability in the Rossmann-fold domain yields split MHT (sMHT) AND gates whose fragments cooperatively associate to synthesize CH_3_Br. Additionally, we identify a sMHT whose fragments require fusion to pairs of interacting partner proteins for maximal activity. We also show that sMHT fragments fused to FKBP12 and the FKBP-rapamycin binding domain of mTOR display significantly enhanced CH_3_Br production in the presence of rapamycin. This gas production is reversed in the presence of the competitive inhibitor of FKBP12/FKPB dimerization, indicating that sMHT is a reversible reporter of post-translational reactions. This sMHT represents the first genetic AND gate that can report on protein-protein interactions via an indicator gas. Because indicator gases can be measured in the headspaces of complex environmental samples, this protein fragment complementation assay should be useful for monitoring the dynamics of diverse molecular interactions within microbes situated in hard-to-image marine and terrestrial matrices.

## Introduction

A wide range of genetically-encoded proteins have been developed for visualizing gene expression in natural and engineered cellular systems, including reporters that produce a pigment^1^, luminescent signal^2^, or fluorescent signature^3^. The utility of these visual reporters has been expanded through the design of a rainbow of colors that can be used in parallel^4^ and the creation of fast biosensors that directly report on post-translational reactions without the need for transcription^5^. One of the earliest reporter successes was a fluorescent indicator for Ca^2+^, which was built by fusing green fluorescent protein (GFP) to calmodulin^6^. Since that time, a wide range of visual reporters have been engineered for real-time sensing of cellular reactions, including protein solubility^7^, protein-protein interactions^8,9^, primary metabolites^10^, secondary metabolites^11^, inorganic cofactors^12^, DNA sequence motifs^13^, and DNA damage^14^. Many of these reporters have been built by creating split proteins whose fragments do not effectively generate an output unless they are fused to pair of proteins that associate and assist with fragment complementations^15^. While these protein-fragment complementation assays (PCAs) can be created using combinatorial design methods^16^, recent studies have shown that they can also be rationally designed using family sequence information^17^.

Existing split protein reporters have proven useful for studying molecular interactions across proteomes in model organisms within laboratory environments, but they have been challenging to apply in opaque and/or highly autofluorescent environments like soils, marine sediments, and wastewater. Techniques for monitoring visual reporters have been developed to enable studies of dynamic processes near the surfaces of high extinction matrices^18^. These studies are performed in containers with windows (*e*.*g*., rhizotrons) that allow for surface imaging of environmental matrices at different depths^19^. These containers have enabled studies that examine a range of environmental processes, such as root discrimination^20^, nutrient bioavailability^21^, and pollutant degradation^22^. While rhizotron studies can provide data about microbial processes near matrix surfaces, they cannot yet report on microbial processes in non-surface soils or marine sediments. In addition, they do not address the problem of measurements in highly autofluorescent environments like seawater.

Recently, a new reporting approach was described that is compatible with non-disruptive detection of gene expression in non-surface environmental matrices such soils and sediments^23^. With this approach, a methyl halide transferase (MHT) is used as a reporter of gene expression. When expressed, MHT synthesizes a rare volatile indicator gas (CH_3_X) using S-adenosylmethionine and halide ions^24^. This gas diffuses out of matrices and can be quantified in the headspace using a gas chromatograph (GC) mass spectrometer (MS)^25^. MHTs have been used as a reporter to monitor how soil conditions influence conjugation^23^, the bioavailability of acylhomoserine lactones that are used for microbial quorum sensing^26^, and the impact of dissolved organic matter on the bioavailability of flavonoids that mediate plant-microbe communication^27^. Although these studies have provided evidence that MHTs are useful for studying how environmental matrices alter the bioavailability of signals that mediate cell-cell communication, gas reporters cannot yet be used to directly monitor post-translational reactions in the same way as visual reporters^6–14,28^.

Members of the DNA methyl transferase family have been split into pairs of fragments that retain the ability to assemble into active enzymes^29,30^. This finding suggests that other members of the methyl transferase superfamily could be engineered to create protein fragment complementation assays^31,32^. While methyltransferases that use DNA and halides as substrates both belong to the class I family, and both contain a conserved Rossmann-like fold, these family members have evolved very different substrate specificities. Additionally, the DNA methyltransferases that were previously split to create a PCA were bisected within their DNA binding regions^29,30^ (**Figure S1**), a region that is not conserved in MHTs. Here, we investigated whether MHT family sequence information could be used to rationally split a MHT within the Rossmann-like fold without disrupting catalytic activity. We find that a split MHT (sMHT) designed using a multiple sequence alignment retains the ability to synthesize CH_3_Br. We show that sMHTs can report on a range of interacting proteins, including pairs of proteins that form stable complexes and pairs of proteins whose interactions are stabilized by the binding of small organic and inorganic compounds. Finally, we demonstrate that sMHT is a reversible reporter of protein-protein interactions and that sMHT activity decreases upon protein complex dissociation.

## Results and Discussion

### Design of a split MHT

Family sequence data has been successfully used to guide the design of protein fragment complementation assays^17^. With this approach, a multiple sequence alignment is used to calculate the positional amino acid sequence variability across a protein family at each native position, a profile illustrating the positional amino acid variability is generated and smoothed using a sliding window, and peptide bonds within regions with moderate to high amino acid sequence variability are targeted for backbone fission. **Figure 1a** shows a profile generated for MHT using 83 family members numbered by the primary structure of *Batis maritima* MHT (*Bm*-MHT). A comparison of this profile with the structure of *Arabidopsis thaliana* MHT (**Figure 1b**) reveals high amino acid variability near the termini^33^. In contrast, residues 40 to 170 contain regions with greater sequence conservation interspersed with small patches of higher sequence variability. The residues with the highest sequence conservation make direct contacts with the S-adenosyl-homocysteine product in the structural model^33^. **Figure 1b** shows that many of the residues that have high variability are distal from the substrate binding pocket. The residues that mediate halide substrate binding are not known.

**Fig. 1.**
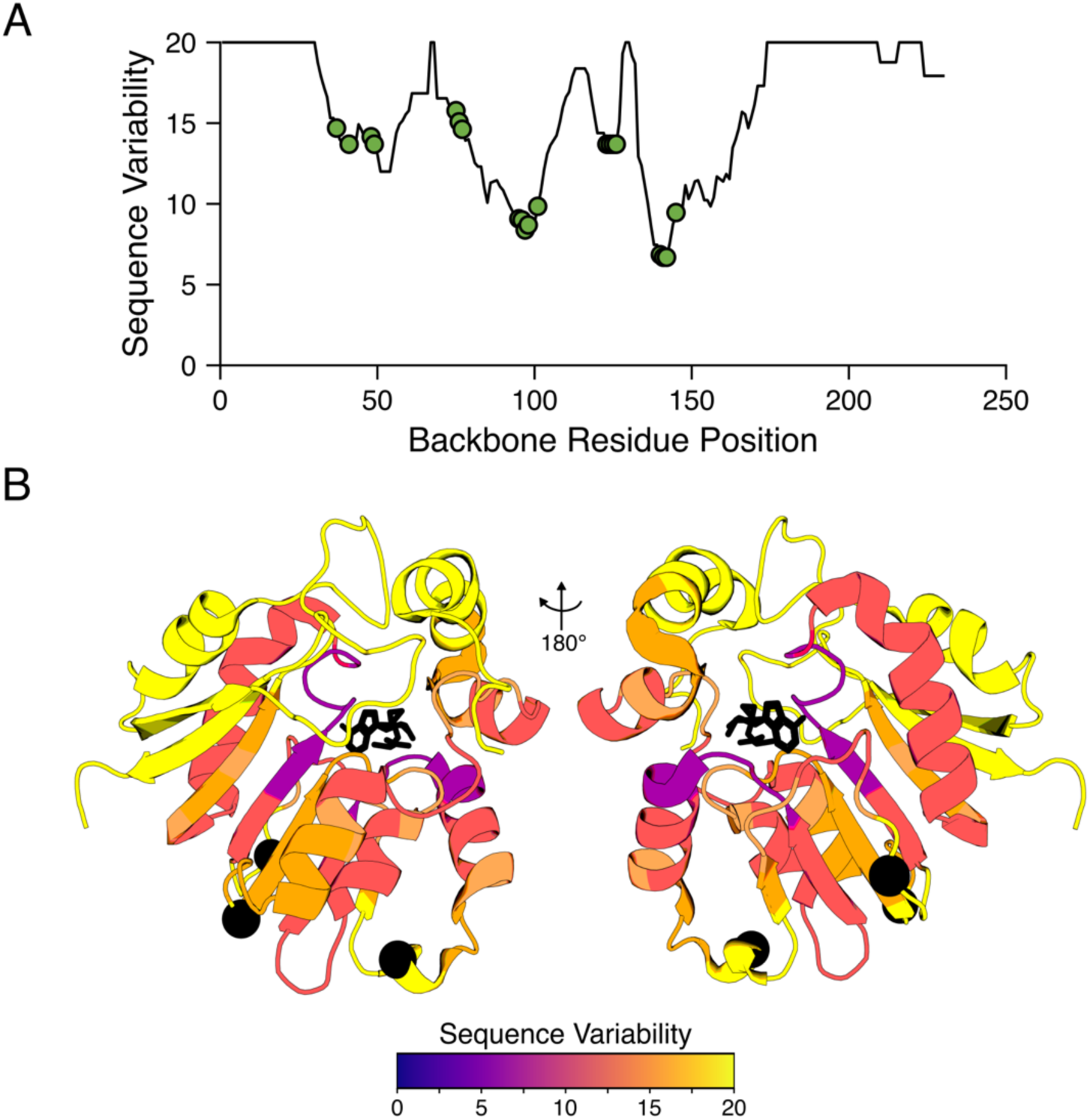
Sequence variability in the MHT family. (**A**) Profile showing the sequence variability in the MHT family. The residues that make contact with the product of the reaction, S-adenosyl-homocysteine^33^, are noted with green circles. (**B**) The product-bound structure of *Arabidopsis thaliana* MHT (PDB: 3LCC) with residues colored to illustrate the sequence variability in the MHT family. Local maxima in sequence variability at residue 67, 113, and 129 are illustrated with black spheres. S-adenosyl-homocysteine is shown as a black stick model.

To investigate if an MHT-fragment complementation assay could be created, we generated three different split MHTs (sMHT) by subjecting *Bm*-MHT to fission at the peptide bonds in the peaks of sequence high variability. We chose this MHT because it has been successfully used as a reporter of gene expression in soils^23,26,27^. Specifically, we targeted the peptide bonds connecting residues 67 and 68 (sMHT-67), 113 and 114 (sMHT-113), and 129 and 130 (sMHT-129). Vectors were initially created that express *Bm-*MHT fragments (F1 and F2) as fusions to SYNZIP17 and SYNZIP18 peptides, respectively (**Figure 2a**). This approach was chosen to build our initial constructs because it has been successfully used to screen for other split proteins that require assistance from interacting proteins to function^17^. The SYNZIP17 and SYNZIP18 synthetic peptides associate strongly (K_d_ = 10 nM) to form an antiparallel coiled-coil^34^. To allow sufficient conformational flexibility for the MHT fragments to associate, they were fused to SYNZIP17 and SYNZIP18 peptides using glycine-rich linkers (GGGGS)_2_AAA.

**Fig. 2.**
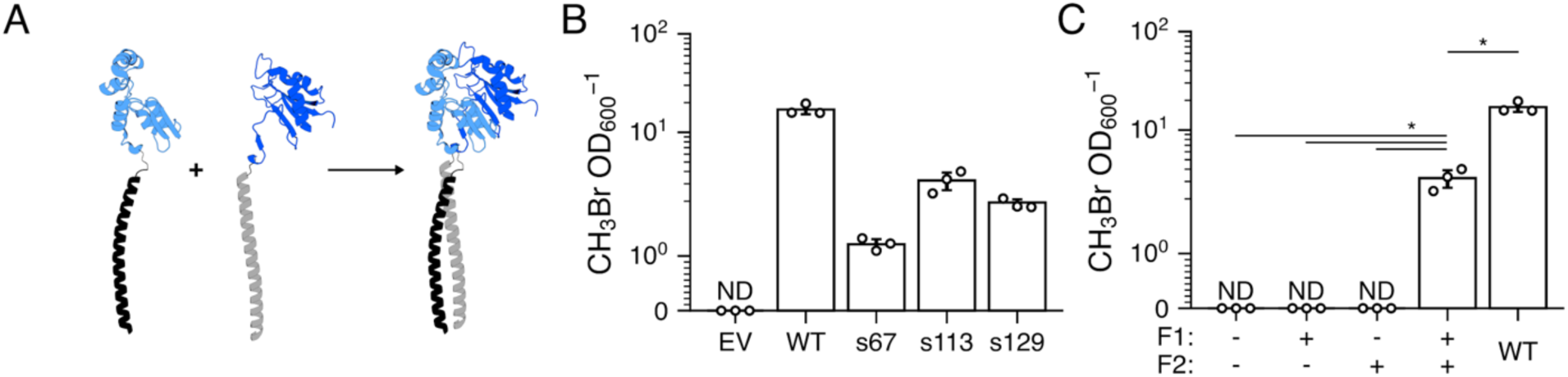
Activity of MHT fragments fused to SYNZIP-17 and SYNZIP-18. (**A**) MHT fragments (light and dark blue) were fused to heterodimerizing SYNZIP coils (light and dark gray) to assist reconstitution of activity. **(B)** CH_3_Br (µM) accumulation in the headspace of cultures after a 24 hour incubation of cells transformed with an empty vector (EV), a vector containing *Bm*-MHT (WT), or vectors with sMHTs fragmented after residues 67 (s67), 113 (s113), or 129 (s129). Relative to cells containing the EV, CH_3_Br accumulation was significantly increased in cells expressing *Bm-*MHT (p=1.75 × 10^−4^), s67 (p=2.35 × 10^−4^), s113 (p=5.11 × 10^−4^), and s129 (p=2.86 × 10^−4^). (**C**) CH_3_Br (µM) accumulation in the headspace of cultures after 24 hour incubations of cells containing a vector that does not express sMHT-113 fragments (-/-) and vectors that express F1-SYNZIP-17 only (+/-), SYNZIP-18-F2 only (-/+), F1-SYNZIP-17 and SYNZIP-18-F2 (+/+), or *Bm*-MHT (WT). The amount of CH_3_Br produced by cells expressing both sMHT-SYNZIP fragments was significantly higher than cells expressing neither fragment, F1-SYNZIP-17 only, or SYNZIP-18-F2 only (p=2.21 × 10^−3^ for each). Cells expressing *Bm-*MHT produced significantly more CH_3_Br than cells expressing F1-SYNZIP-17 and SYNZIP-18-F2 (p=5.11×10^−4^). Each data point represents the average of 3 biological replicates, with error bars representing +/- 1 standard deviation. Individual replicates are shown as circles. Significance differences (p < 0.01) noted with an asterisk were calculated using a two-sided, independent t-test. ND = non-detectable.

### Multiple split MHT synthesize CH_3_Br

To determine which sMHT produce the largest signal when expressed in cells, we examined the level of indicator gas (CH_3_Br) that accumulated in the headspace of *Escherichia coli* cultures coexpressing the fragments of the different sMHTs fused to the SYNZIP peptides (F1-SYNZIP17 and SYNZIP18-F2). As a frame of reference, we evaluated the level of CH_3_Br synthesized by cells expressing *Bm*-MHT or containing an empty vector. We chose to monitor CH_3_Br production because *Escherichia coli* lacking an MHT does not synthesize detectable levels of this volatile chemical even when grown in 100 mM NaBr (**Figure 2b**). After a 24-hour incubation in closed vials, all cultures expressing a native or engineered MHT accumulated measurable CH_3_Br. In addition, all three of the sMHT fusions produced significantly (p < 0.01) higher CH_3_Br than cells containing an empty vector, which did not produce detectable CH_3_Br. All of the sMHT produced ≥5-fold less CH_3_Br than the *Bm-*MHT. However, sMHT-113 produced the most CH_3_Br of any of the sMHTs, 2.7-fold more than sMHT-67 and 1.6-fold more than sMHT-129.

We chose to focus all subsequent analysis on sMHT-113, since this pair of fragments presented the strongest whole cell activity. To determine if sMHT-113 requires both fragments to function, we created vectors that express the sMHT-SYNZIP fragments individually and compared the gas produced by cells expressing both fragments, the individual fragments, and full length *Bm-*MHT. Cells expressing the individual sMHT-113 fragments did not produce any detectable CH_3_Br, similar to cells containing an empty vector (**Figure 2c**).

In a recent study, we found that *Bm*-MHT activity in *E. coli* increases as growth temperature is increased from 30 to 37°C^26^. To investigate how temperature affects sMHT-113 activity, we evaluated the indicator gas produced by cells coexpressing sMHT-113 fragments across the same temperature range (**Figure S2**). For these experiments, we measured how the rate of gas production changed over six hours by monitoring the gas accumulation each hour and normalizing those values to OD_600_. The gas production rate at 37°C was significantly lower than that observed at 30°C. This observation can be contrasted with a previous study that found that *Bm*-MHT presents activity at 37°C that is 2.5-fold higher than that observed at 30°C^26^. These differences are interpreted in arising because backbone fragmentation decreases MHT stability, lowering the temperature of maximal activity.

### Using a split MHT as a genetic AND gate

To investigate whether sMHT-113 can be used as an AND gate to report on the activity of a pair of conditional promoters, like a split transcription factor^35^, we built a construct that expresses F1-SYNZIP17 and SYNZIP18-F2 as fusions to GFP and red fluorescent protein (RFP), respectively (**Figure 3a**). The anhydrotetracycline (aTc) responsive transcription factor TetR was used to regulate F1-SYNZIP17-GFP expression, while the isopropyl β-D-1-thiogalactopyranoside (IPTG) responsive transcription factor LacI was used to control RFP-SYNZIP18-F2 expression. Using *E. coli* XL1 transformed with this construct, we evaluated the effects of different combinations of inducer concentrations on CH_3_Br production and whole cell fluorescence following a 24-hour incubation. In these experiments, cells grew to similar optical densities across the treatments (**Figure S3**). Consistent with AND gate behavior, both aTc and IPTG were required to obtain the largest gas signal (**Figure 3b**). In the absence of aTc, the CH_3_Br production was <1% of the maximum level over the full range of IPTG concentrations (**Figure 3b**). In the absence of IPTG, the CH_3_Br production was undetectable at the two lowest aTc concentrations evaluated, although it was 20% of the maximum level observed at the highest aTc concentration. The gas production observed at the highest aTc concentration and in the absence of IPTG was interpreted as arising from IPTG-inducible promoter leakiness, which has been previously observed^35,36^.

**Fig. 3.**
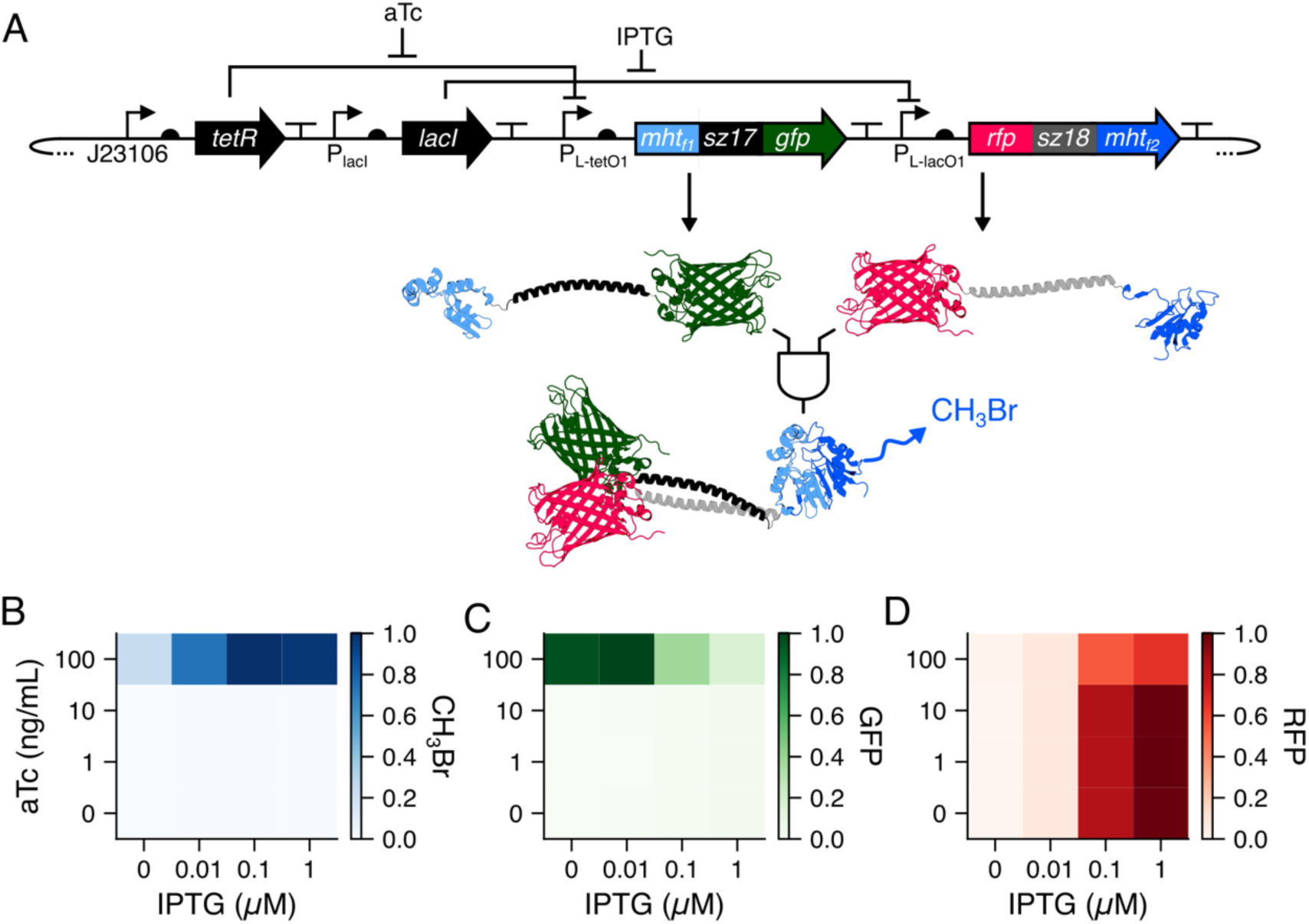
Using the split MHT as a two-input AND gate. (**A**) The genetic circuit used to regulate transcription of the genes encoding the MHT fragments. Expression of F1-SYNZIP-17-GFP and RFP-SYNZIP-18-F2 was induced with aTc and IPTG, respectively. (**B**) Relative CH_3_Br production by cultures following a 24-hour incubation in the presence of different aTc (0, 1, 10, 100 ng/mL) and IPTG (0, 0.01, 0.1, and 1 mM) concentrations. The relative CH_3_Br production was calculated as the ratio of CH_3_Br concentration to OD_600_ to account for variability in cell growth, and these values were scaled from 0 to 1. CH_3_Br production was significantly higher in the presence of 100 ng/mL aTc and 1 mM IPTG than in experiments containing no inducer (p = 2.96 × 10^−4^), 100 ng/mL aTc alone (p = 9.68 × 10^−4^), or 1 mM IPTG alone (p = 3.04 × 10^−4^). Relative (**C**) green and (**D**) red fluorescence of cells following CH_3_Br measurements. The relative fluorescence was calculated as the ratio of emission to OD_600_ to account for variability in cell growth, and these values were scaled from 0 to 1. Each data point represents the average of 3 biological replicates. All p-values were calculated using a two-sided, independent t-test.

Fluorescence measurements also revealed sMHT fragment expression consistent with AND gate behavior. The addition of aTc alone increased the GFP signal (**Figure 3c**), while it had little effect on the RFP signal (**Figure 3d**). In addition, the RFP signal increased with IPTG, with little effect on GFP across 0-10 ng/mL aTc. At 100 ng/mL aTc, the decrease in GFP signal with increasing IPTG concentration is interpreted as arising from FRET between GFP and RFP when both fluorescent proteins are highly expressed^37^. Taken together, these indicator gas and fluorescence measurements provide evidence that our sMHT requires two fragments to function. Furthermore, these results illustrate how this protein can be used as a genetic AND gate to report on multiple environmental conditions.

### Reporting on different protein-protein interactions

To directly investigate whether sMHT-113 requires assistance from a protein-protein interaction for maximal catalytic activity, we compared the gas production of cells expressing F1-SYNZIP17 and SYNZIP18-F2 with cells expressing F1 and SYNZIP18-F2. The fragments in both of these sMHT are expected to present similar translation initiation rates when expressed from the same ribosomal binding site (RBS), because removal of the gene encoding SYNZIP17 does not affect the genetic context of the RBS used for translation initiation^38^. **Figure 4a** shows that *E. coli* expressing F1-SYNZIP17 and SYNZIP18-F2 yield significantly higher gas production than cells expressing F1 and SYNZIP18-F2 at 30°C. The ratio of gas production between cells expressing F1 and SYNZIP18-F2 versus F1-SYNZIP17 and SYNZIP18-F2 is temperature-dependent. Cells expressing F1-SYNZIP17 and SYNZIP18-F2 produced 49.8-fold more CH_3_Br at 25°C than those expressing F1 and SYNZIP18-F2, 26.6-fold more CH_3_Br at 30°C, and 20.9-fold more CH_3_Br at 37°C (**Figure S4a**).

**Fig. 4.**
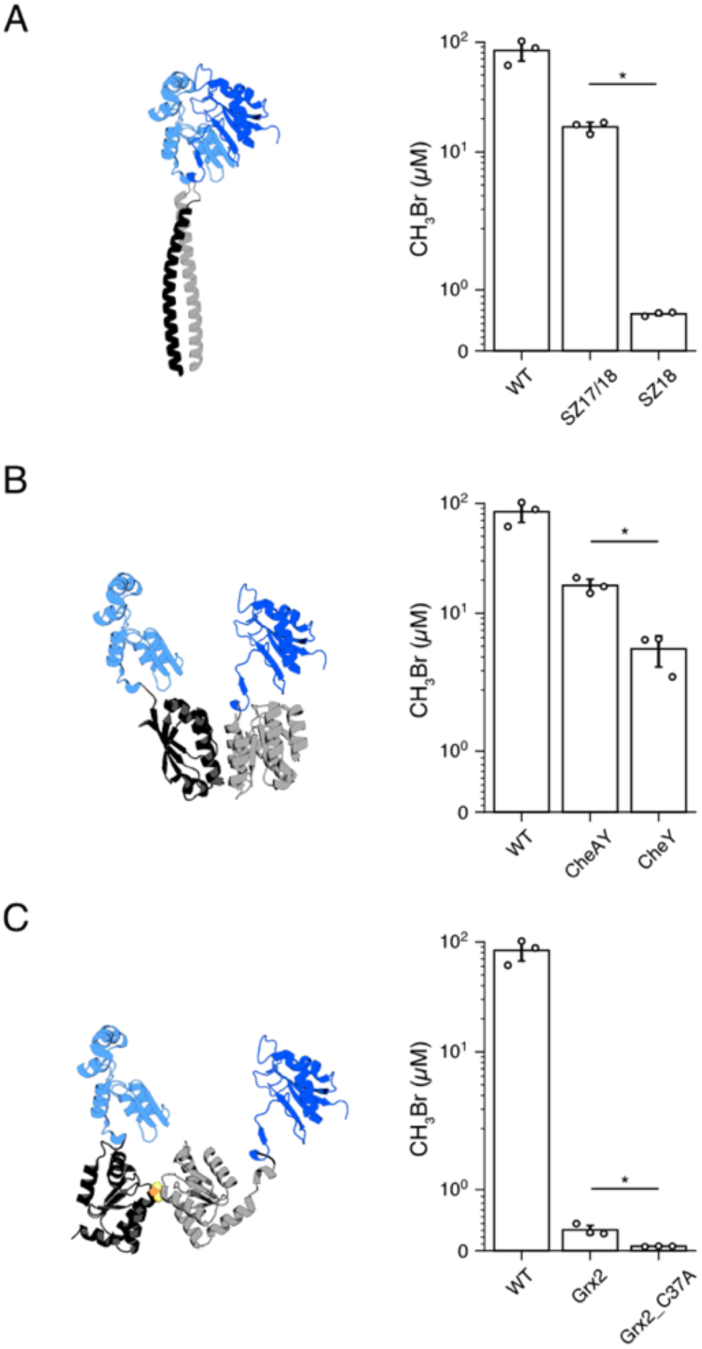
Split MHT activity is enhanced by different pairs of interacting proteins. To evaluate the effect of different pairs of interacting proteins on MHT activity, we compared the CH_3_Br synthesized by cells coexpressing sMHT-113 F1 and F2 fused to three pairs of dimerizing proteins, including (**A**) SYNZIP-17 and SYNZIP-18 (SZ17/18), which produced significantly (p = 1.95 × 10^−4^) more CH_3_Br than cells expressing F1 alone and F2-SYNZIP-18 (SZ18), (**B**) F1-CheA and CheY-F2 (CheAY), which produced significantly (p = 2.77 × 10^−3^) more CH_3_Br than cells expressing F1 and F2-CheY (CheY), and (**C)** F1-GRX2 and F2-GRX2 (Grx2), which produced significantly (p = 6.63 × 10^−3^) more CH_3_Br than cells expressing Grx2 C37A mutants that cannot coordinate the iron-sulfur cluster necessary for dimerization (Grx2_C37A). *Bm-*MHT (WT) is shown for comparison. Bars represent the average of 3 biological replicates, with error bars representing +/- 1 standard deviation. Individual replicates are shown as circles Significance differences (p < 0.01) noted with an asterisk were calculated using a two-sided, independent t-test.

We next investigated the modularity of sMHT-113 by testing if a pair of globular proteins that form a stable complex could also be used to enhance the activity of sMHT. To test this idea, we fused F1 and F2 to CheY and the CheY-binding domain of CheA, a pair of heterodimerizing *Thermotoga maritima* chemotaxis proteins^39^. These chemotaxis proteins display high affinity for one another (K_d_ = 200 nM)^39^, like SYNZIP17 and SYNZIP18. However, the termini that are covalently attached to the sMHT fragments are more distant from one another (30 Å) compared with those in SYNZIP17 and SYNZIP18 (<10 Å)^34,40^. When cells expressed sMHT-113 fragments as fusions to CheA and CheY (F1-CheA and CheY-F2), similar levels of CH_3_Br were observed as with cells expressing the SYNZIP17/18 fusions (**Figure 4b**). Additionally, cells expressing F1-CheA and CheY-F2 presented significantly higher gas production compared with cells expressing F1 and CheY-F2. The ratios of gas production with fragment fusions containing or lacking CheA were similar across all temperatures analyzed, with gains of 6.7 at 25°C, 3.8 at 30°C, and 6.3 at 37°C (**Figure S4b**).

To determine if CH_3_Br gas production can be controlled using a protein-protein interaction with even larger separation of the fused termini, we analyzed gas produced by *E. coli* expressing sMHT-113 fragments that were fused to human glutaredoxin (GRX2). Previous studies have shown that GRX2 homodimerizes through an 2Fe-2S cluster^41^. The 2Fe-2S cluster is required for dimerization because GRX2 uses the active site cysteine as ligands for the iron-sulfur cluster^42^. Structural studies have identified Cys37 as the iron ligand in GRX2, and have also revealed that the N- and C-termini in the GRX2 homodimer bridged by a 2Fe-2S cluster are separated by >57 Å^42^. When cells expressed the sMHT-113 fragments as fusions to GRX2 (F1-GRX2 and GRX2-F2), CH_3_Br production was detected across all temperatures tested (**Figure 4c**). However, this gas production was more than an order of magnitude lower than gas production by the SYNZIP17/18 fusions at all temperatures tested (**Figure S4c**). In part, this lower signal arises because F1-GRX2 and GRX2-F2 can homodimerize to form complexes that do not produce gas.

To evaluate whether F1-GRX2 and GRX2-F2 require a bridging 2Fe-2S for maximal gas production, we mutated Cys37 to an Ala in both fragment fusions and tested gas production under identical conditions. At all temperatures, indicator gas production by F1-GRX2 and GRX2-F2 was significantly higher than the C37A mutants. The ratios of gas production by GRX2/GRX2-C37A was 9.4 at 25°C, 5.5 at 30°C, and 1.5 at 37°C. Taken together, our results from the SYNZIP17/18, CheA/Y, and GRX/GRX2 fusions suggest that sMHT can report on the formation of protein complexes with both proximal and distal termini.

### Monitoring chemical complementation using a split MHT

Split proteins have been previously used to report on chemicals by fusing their fragments to a pair of proteins whose interaction is stabilized by chemical binding (**Figure 5a**)^43,44^. One chemical-dependent protein-protein interaction that has been widely used to create such systems is the FK506-binding protein (FKBP) and FKBP-rapamycin binding domain of mTOR (FRB)^8,11,17,43,45^. In the crystal structure of FKBP complexed with FRB and rapamycin^46^, the N- and C-termini of these proteins are ∼34 Å apart. Since this distance is similar to the separation of the termini in the CheA/Y complex^39^, which yielded an indicator gas signal when fused to sMHT, we hypothesized that the activity of sMHT-113 could be controlled by rapamycin if we fused the fragments to FKBP and FRB. To test this idea, we created a construct that express F1-FKBP and FRB-F2 and evaluated CH_3_Br production in the presence and absence of rapamycin. Because protein expression in this construct is repressed by LacI, we first analyzed the effects of IPTG and rapamycin on CH_3_Br production (**Figure S5**). We then evaluated the effects of a range of rapamycin concentrations on gas production in the presence of a constant IPTG concentration (**Figure 5b**). Addition of rapamycin led to significant increases in CH_3_Br production compared with untreated cells, with the most significant gain in CH_3_Br production (18.5-fold) observed with 25 µM rapamycin.

**Fig. 5.**
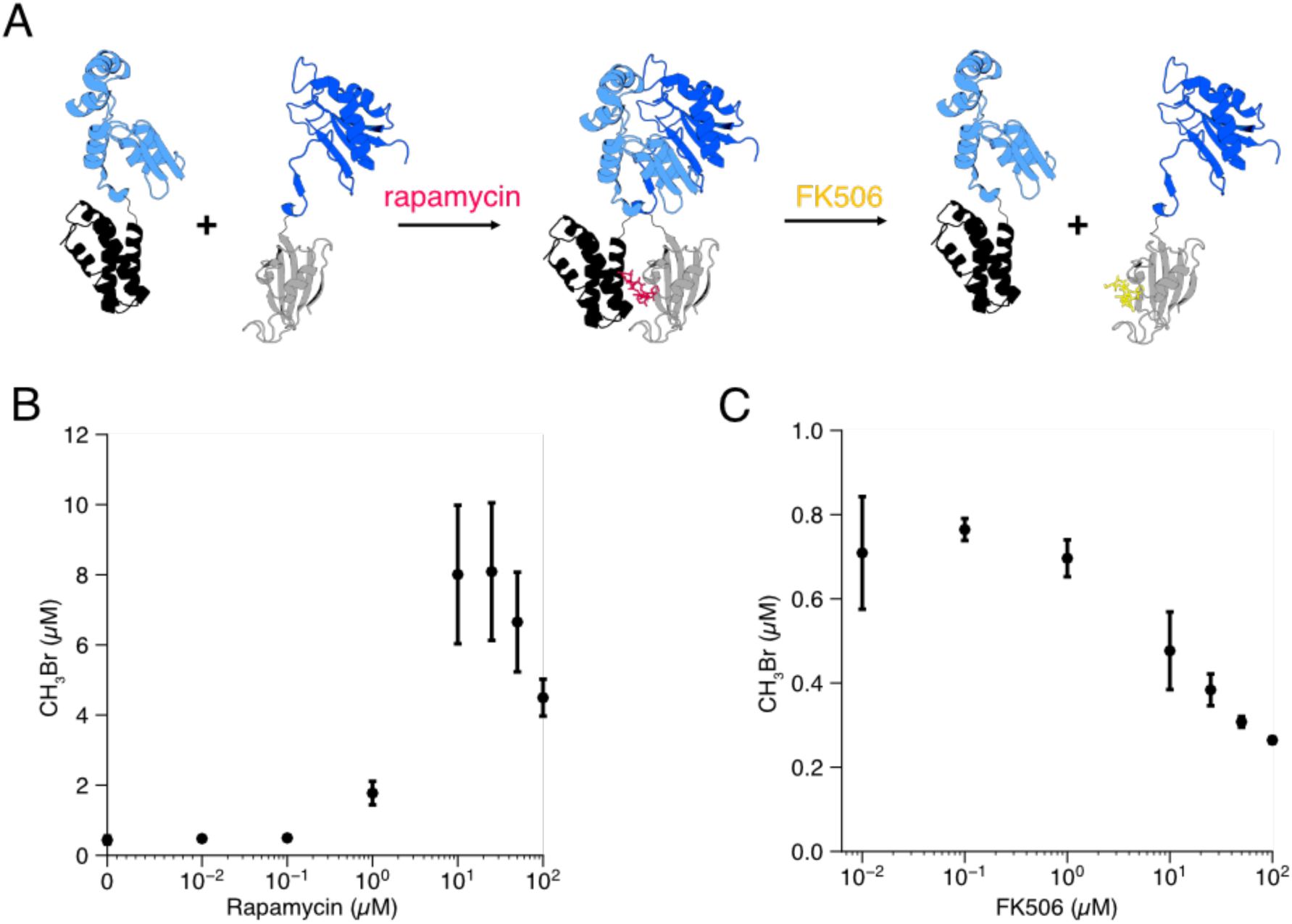
Using the split MHT to monitor chemical-induced dimerization. (**A**) Rapamycin induces F1-FKBP and FRB-F2 dimerization, while FK506 is a competitive inhibitor of rapamycin that inhibits dimerization following F1-FKBP and FRB-F2 dissociation. (**B**) The effect of different rapamycin concentrations on CH_3_Br production by cells expressing F1-FKBP and FRB-F2. The CH_3_Br accumulation observed with 25 µM rapamycin is significantly greater than the accumulation in cultures lacking rapamycin (p = 5.3 × 10^−3^). (**C**) To evaluate sMHT reversibility, cultures containing inducer (1 mM IPTG) and rapamycin (10 µM) cells were grown for 3 hours, cells were washed and resuspended in medium containing varying concentrations of FK506, and gas production was measured following incubation of cultures at 37°C for 24 hours. Data represent the average of 3 biological replicates, with error bars representing +/- 1 standard deviation.

Some split proteins form irreversible complexes, such as split fluorescent proteins^47^. To investigate if the association of sMHT-113 fragments is reversible, we evaluated the effect of adding FK506, a competitive inhibitor of rapamycin binding, on indicator gas production^48,49^. For these experiments, exponential phase cultures of *E. coli* expressing F1-FKBP and FRB-F2 were grown in the presence of rapamycin (10 µM) and IPTG (1 mM) for three hours. At this point, the cells were washed with M63 medium to remove the rapamycin. Cells were then resuspended in fresh medium containing varying concentrations of FK506 but lacking IPTG, which switches off F1-FKBP and FRB-F2 expression, and incubated overnight prior to measuring CH_3_Br (**Figure 5c**). Gas measurements from this experiment reveal a pattern of CH_3_Br production inversely proportional to the concentration of FK506. This finding suggests that sMHT-113 is a reversible reporter of protein-protein interactions. If sMHT-113 association was irreversible, the gas signal would have remained similar across all FK506 concentrations. **Implications for synthetic biology**. The sMHT fragments characterized herein presented enhanced CH_3_Br synthesis rates when the fragments were fused to different pairs of interacting proteins, indicating that the sMHT-113 functions as a PCA^15^. Because sMHT generates an indicator gas as an output, this new PCA will enable detection of protein-protein interactions under hard-to-image conditions (*e*.*g*., in marine sediments, soils, and wastewater)^23,26^ where other PCAs cannot be used due to their generation of a visual signal^6–14,28^. Additionally, the sMHT AND gate described herein will be a useful synthetic biology tool for environmental studies that probe micron-scale spatial heterogeneity in hard-to-image matrices. For example, this AND gate could report on the bioavailability of a metals in the environment AND the expression of metal-dependent oxidoreductases that drive biogeochemical cycling or greenhouse gas production in order to determine whether the oxidoreductases have access to metallic cofactors required for catalysis. Finally, the observation that sMHT-113 functions when fused to a variety of interacting proteins whose termini have a wide range of proximities (∼10 to 60 Å) suggests that this sMHT will be useful as a readout for a diverse set of natural and synthetic protein interactions.

Like other PCAs^50^, sMHT-113 presented a small background signal when fragments were fused to only one protein. Because previous studies have shown that *E. coli* does not produce detectable CH_3_Br unless it expresses a non-native MHT gene^23^, this small signal is interpreted as arising from sMHT fragments associating without assistance. Biochemical studies examining how protein fragment structure influences split protein function have suggested that residual structure in the fragments may influence such background signals^51^. A study examining the cooperative function of a split adenylate kinase found that fragments derived from a thermostable family member retained more residual structure and exhibited higher activity than identical fragments from a less stable homolog^51^. This finding also suggests that one way to decrease the background of our MHT-fragment complementation assay in the future will be to split homologs with lower thermostability or incorporate mutations that destabilize residual structure.

In the future, it will be interesting to explore the diversity of conditional protein-protein interactions that can be monitored using sMHT-113 and whether this split protein can be coupled to commonly used sensing systems in synthetic biology. For example, many two-component sensing systems^52^ include response regulators that that exhibit enhanced association when the sensing system is activated^53^. This change in response regulator association can be challenging to monitor directly *in vivo* using visual FRET-type sensing. Because sMHT produces a signal that arises from catalysis and has effectively no background in some microbial systems^23^, this PCA may be more sensitive than more traditional visual reporting strategies. Overall, we expect sMHT to be a useful and sensitive tool to study a broad range of protein-protein interactions in both ideal laboratory environments and hard-to-image environmental matrices.

## Methods

### Materials

Antibiotics were from Research Products International and Sigma-Aldrich, rapamycin and FK506 were from Tokyo Chemical Industry, and all other chemicals were from Sigma-Aldrich, VWR, or BD Biosciences. DNA was from Integrated DNA Technologies, enzymes were from New England Biolabs and Thermo Fisher Scientific, and kits for DNA purification were from Zymo Research and Qiagen. Vials for gas measurement were from Phenomenex. *Escherichia coli* XL1 was used for all DNA manipulations and analysis of fluorescent protein fusions, while gas reporting was performed in *E. coli* MG1655, *E. coli* XL1, and *E. coli* CS50^36^.

### Sequence alignment of MHT homologs

A multiple sequence alignment was generated using the sequences of 83 MHTs that have been heterologously expressed as active enzymes in *E. coli*^54^ using the MUSCLE algorithm^55^. Positional amino acid variability was calculated as the number of unique amino acids observed at each native site. Any sites containing a gap in one MHT sequence were given a value of 20. At each position, we report the positional variability as the average of a sliding window of fourteen residues, which is numbered using the *Bm-*MHT sequence. Visualization of sequence variability on tertiary structure was done using the PyMol molecular visualization system (github.com/schrodinger/pymol-open-source) and residues were colored using pymolcolorizer (github.com/jatk/pymolcolorizer).

### Growth medium

All growth for molecular biology was performed in Lysogeny Broth (LB), while studies analyzing CH_3_Br production used a modified M63 medium containing 100 mM NaBr^26^. This medium yields a high indicator gas signal with *E. coli* expressing *Bm*-MHT without affecting cell fitness^23^.

### Plasmids

All plasmids are listed in **Table S1**. Plasmids were assembled using Golden Gate assembly of amplicons generated using PCR^56^, and all vectors were sequence verified. As templates for vector backbones, we used either (1) pSR353 (pEMF plasmids)^57^, (2) pET-28b (pDH001 – 011 and pDH013), or (3) pSAC01^17^ (pDH012). As templates for creating inserts, we used vectors containing the *Bm-*MHT gene^23^, GFPmut3^58^ and mRFP1^59^ genes, and DNA encoding pairs of interacting proteins with a kanamycin resistance marker (*kan*^*R*^), including *synzip-kan*^*R*^ (SYNZIP17/18), *sz18-kan*^*R*^ (SYNZIP18 only), *ay-kan*^*R*^ (CheA/Y), *y-kanR* (CheY only)^17,35,45,50,60^. Vectors containing these DNA sequences were cloned with regulated promoters and the sMHT fragments, such that the sMHT fragments were expressed as fusions to the interacting proteins. A DNA cassette (*grx-grx-kan*^*R*^) encoding a pair of human glutaredoxin 2 (GRX2) was created by synthesizing two copies of the GRX2 gene (each with a different coding sequence) and cloning these genes into a vector containing the same regulatory elements used as the other sMHT-protein fusions. A mutant of this cassette (*c37a-c37a-kan*^*R*^) was created by mutating the active site cysteines (C37A) in both GRX2 genes. GRX2-C37A mutants lack both cysteines used as iron ligands^12^, and thus cannot coordinate a 2Fe-2S cluster. To regulate expression, we used the TetR-repressed P_Ltet-O1_ promoter, which can be derepressed by anhydrotetracycline (aTc)^61^, the LacI-repressed P_Llac-O1_ promoter, which can be derepressed by isopropyl β-D-1-thiogalactopyranoside (IPTG)^61^, or the LacI-regulated T7 promoter^62^, which can also be derepressed by IPTG. All vectors used synthetic RBS, which were designed using a thermodynamic model^38^.

### Indicator gas measurements

To assess the activity of the sMHT variants fused SYNZIP coils, we measured CH_3_Br production by *E. coli* MG1655 transformed with the different constructs. For this experiment, overnight cultures in LB with chloramphenicol (34 µg/mL) were diluted 1:100 into M63 medium with the same antibiotic and grown to mid-log phase (OD_600_ = 0.6-0.8) at 30°C, while shaking at 250 rpm. aTc (100 ng/mL) was then added to induce sMHT-SYNZIP production, and 1 mL of each culture was transferred to a 2 mL VEREX™ glass vial. Vials were capped with VEREX™ crimp tops (11mm diameter, PTFE/Sil, silver) and incubated for at 30°C while shaking at 250 rpm. After 24 hours of incubation, headspace gas was analyzed using GC-MS. To assess the activity of sMHT-113 fused to different interacting proteins, the fusions were expressed in *E. coli* CS50. In these experiments, protein expression was induced with IPTG (1 mM) and cultures were incubated at 30°C for 18 hours before measuring CH_3_Br accumulation.

Gas production was analyzed using either (1) an Agilent 7890B gas chromatograph (GC) connected to an Agilent 5779E mass spectrometer (MS) via a DB-VRX capillary column (20 m, 0.18 mm ID, 1 µm film), or (2) an Agilent 7820A GC connected to a 5779B MS via a PoraPLOT Q capillary column (24 m, 0.25 mm ID, 8 µm film). For both GC-MS setups, vials were loaded with an Agilent 7693A autosampler and headspace gas (50 µL) was injected using a 100 µL gastight syringe (Agilent G4513-80222). In the first GC-MS setup, the oven temperature started at 45°C and held at 45°C for 84 seconds, then ramped at 36°C per minute to 60°C and held at 60°C for 9 seconds. In the second GC-MS setup, the oven temperature started at 85°C and ramped at 12°C per minute to 105°C then ramped at 65°C per minute until 150°C and holding for 144 seconds at 150°C. In both GC-MS setups, the MS was configured for selected ion monitoring mode for methyl bromide isotopes (MW = 93.9 and 95.9 for CH_3_^79^Br and CH_3_^81^Br, respectively). Agilent MassHunter WorkStation Quantitative Analysis software was used to quantify the relative peak area for each sample. The peak area of the major isotope (CH_3_^79^Br) was used to quantify the accumulated gas per sample while the minor isotope (CH_3_^81^Br) was used as a qualifier. A standard curve for relating the peak area to the amount of CH_3_Br was generated using serial dilutions of bromomethane (100 µg/mL in methanol, Sigma-Aldrich or 2000 µg/mL in methanol, Restek) in M63 media. After each gas measurement, vials were uncapped, and the density of each culture (OD_600_) was measured using a Varian Cary 50 UV-visible spectrophotometer or a DeNovix DS-11 series spectrophotometer.

### Fluorescence spectroscopy

*E. coli* XL1 transformed with the vector that expresses sMHT-113 fragments fused to GFP and RFP were analyzed for CH_3_Br production as described above. Cultures were grown for 24 hours at 30°C in capped vials containing chloramphenicol (34 µg/mL) and different concentrations of IPTG and aTc. Following gas measurements, each culture was pelleted and resuspended in an equal volume of phosphate buffered saline. Following incubation at 37°C for 1 hour to allow for fluorophore maturation^63^, samples were transferred on a 96 well plate (Corning #3603), and optical density (600 nm) and fluorescence were analyzed using a M1000 Pro plate reader (Tecan). For GFP analysis, λ_ex_ = 501 nm and λ_em_ = 515 nm were used. For RFP measurements, λ_ex_ = 550 nm and λ_em_ = 611 nm were used.

### Chemical-induced dimerization experiments

To evaluate the effects of rapamycin on gas production, *E. coli* CS50 harboring a construct containing sMHT-FKBP/FRB was grown to stationary phase in M63 medium containing kanamycin (50 µg/mL) while shaking at 250 rpm. These cultures were diluted 1:100 into M63 medium with the same antibiotic and grown to an OD ∼0.5 at 37°C while shaking at 250 rpm. To induce protein expression, IPTG (1 mM) was added to 1 mL cultures in VEREX™ vials. Varying concentrations of rapamycin were then added, vials were capped, and cultures were incubated at 37°C with 250 rpm shaking. After 24 hours of incubation, gas production was quantified using GC-MS. To investigate whether the rapamycin-induced gas production is reversible, *E. coli* CS50 transformed with the sMHT-FKBP/FRB construct was grown to mid log phase (OD ∼0.5) at 37°C in the presence of IPTG (1 mM) and rapamycin (10 µM). The culture (10 mL) was then incubated aerobically at 37 °C for three hours before the cells were pelleted and washed twice with M63 medium containing kanamycin (50 µg/mL). The cell pellets were resuspended in M63 medium, 1 mL aliquots were mixed with varying concentrations of FK506 and kanamycin (50 µg/mL), and the culture was incubated in capped vials at 37 °C while shaking at 250 rpm. Following 24 hours of incubation, indicator gas was measured using GC-MS.

### Statistics

Error bars in all figures represent standard deviations from three independent biological replicates. P-values were calculated using two-sided, independent t-tests.

## Acknowledgements

We are grateful for financial support from the W. M. Keck Foundation, Moore Foundation, Defense Advanced Research Projects Agency (HR0011-19-2-0019), and National Science Foundation (1150138). JTA was supported by National Science Foundation Graduate Research Fellowships and a Lodieska Stockbridge Vaughn Fellowship, and ML was supported by a Beckman Scholars Fellowship.

