## Supplementary Materials for "A split methyl halide transferase AND gate that reports by synthesizing an indicator gas"

**Table S1.** Constructs used in this study. sMHT(1-x) and sMHT(y-230) indicate the amino acids of *Bm*-MHT included in each sMHT fragment. Sequences will be made available through Addgene, but have not yet been deposited.

| Plasmid | Description | ORF1 | | ORF2 | | Addgene ID |
|---|---|---|---|---|---|---|
| | | Promoter (Regulation) | ORF | Promoter (Regulation) | ORF | |
| pEMF034 | sMHT-113 | $P_{LtetO-1}$ (TetR) | sMHT(1-113)-SYNZIP17 | $P_{LtetO-1}$ (TetR) | SYNZIP18-sMHT(114-230) | not yet deposited |
| pEMF035 | sMHT-67 | $P_{LtetO-1}$ (TetR) | sMHT(1-67)-SYNZIP17 | $P_{LtetO-1}$ (TetR) | SYNZIP18-sMHT(68-230) | not yet deposited |
| pEMF036 | sMHT-129 | $P_{LtetO-1}$ (TetR) | sMHT(1-129)-SYNZIP17 | $P_{LtetO-1}$ (TetR) | SYNZIP18-sMHT(130-230) | not yet deposited |
| pEMF037 | Wild type (WT) control | $P_{LtetO-1}$ (TetR) | *Bm*-MHT | - | - | not yet deposited |
| pEMF038 | Empty vector (EV) control | $P_{LtetO-1}$ (TetR) | - | - | - | not yet deposited |
| pEMF047 | sMHT-113 F1 only | $P_{LtetO-1}$ (TetR) | sMHT(1-113)-SYNZIP17 | - | - | not yet deposited |
| pEMF044 | sMHT-113 F2 only | $P_{LtetO-1}$ (TetR) | SYNZIP18-sMHT(114-230) | - | - | not yet deposited |
| pDH013 | sMHT-113 GFP/RFP | $P_{LtetO-1}$ (TetR) | sMHT(1-113)-SYNZIP17-GFPmut3 | $P_{LlacO-1}$ (LacI) | mRFP1-SYNZIP18-sMHT(114-230) | not yet deposited |
| pDH001 | $P_{T7}$-*Bm*-MHT | T7 (LacI) | *Bm*-MHT | - | - | not yet deposited |
| pDH003 | $P_{T7}$-sMHT-113 | T7 (LacI) | sMHT(1-113)-SYNZIP17 | T7 (LacI) | SYNZIP18-sMHT(114-230) | not yet deposited |
| pDH006 | sMHT-113 SYNZIP18 only | T7 (LacI) | sMHT(1-113) | T7 (LacI) | SYNZIP18-sMHT(114-230) | not yet deposited |
| pDH007 | sMHT-113 CheAY | T7 (LacI) | sMHT(1-113)-CheA | T7 (LacI) | CheY-sMHT(114-230) | not yet deposited |
| pDH008 | sMHT-113 CheY only | T7 (LacI) | sMHT(1-113) | T7 (LacI) | CheY-sMHT(114-230) | not yet deposited |
| pDH009 | sMHT-113 GRX2 | T7 (LacI) | sMHT(1-113)-GRX2 | T7 (LacI) | sMHT(114-230)-GRX2 | not yet deposited |
| pDH010 | sMHT-113 GRX2_C37A mutant | T7 (LacI) | sMHT(1-113)-GRX2_C37A | T7 (LacI) | sMHT(114-230)-GRX2_C37A | not yet deposited |
| pDH011 | sMHT-113 FKBP/FRB | T7 (LacI) | sMHT(1-113)-FKBP | T7 (LacI) | FRB-sMHT(114-230) | not yet deposited |
| pDH012 | P14-sMHT-113 | P14[1] (constitutive) | sMHT(1-113)-SYNZIP17 | P14[1] (constitutive) | SYNZIP18-sMHT(114-230) | not yet deposited |

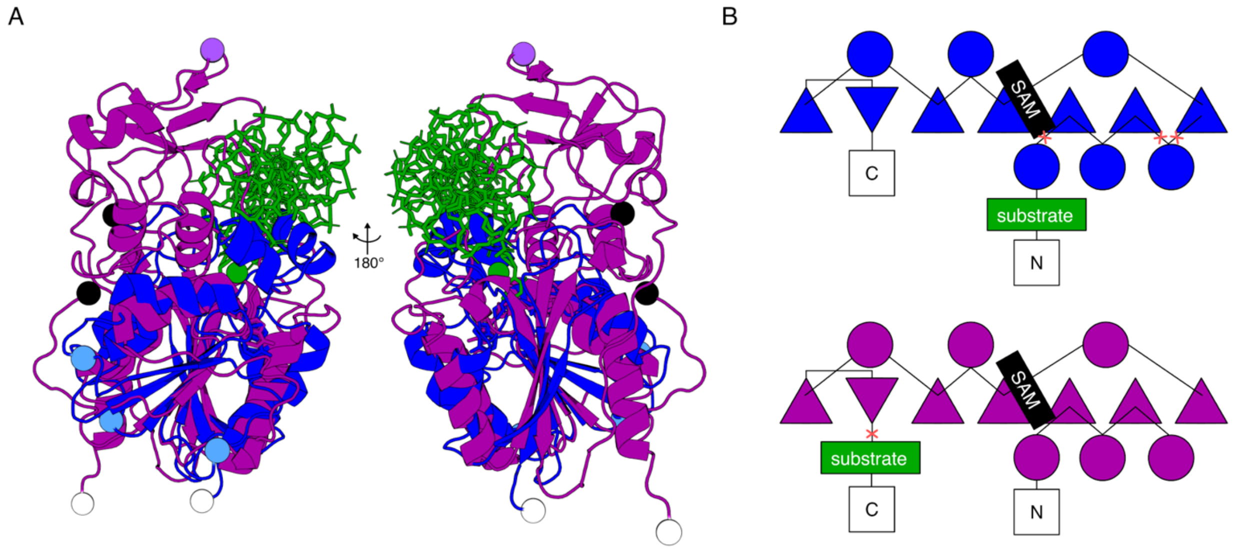


**Figure S1. Comparison of methyl-halide transferase to C5-DNA methyltransferase. (A)** Comparison of tertiary structures of the methyl halide transferase from *Arabidopsis thaliana At*-MHT (blue) and the C5-DNA methyltransferase from *Haemophilus haemolyticus* M.HhaI (purple). Structures of *At*-MHT (PDB: 3LCC)[2] and M.HhaI (PDB: 1MHT)[3] were aligned using the MATT structural alignment algorithm[4] revealing a core region of 138 residues and a RMSD of 3.317 Å. The proposed halide-ion substrate position in the active site of *At*-MHT (green sphere)[2]. The cytosine from the DNA substrate of M.HhaI (green sticks) can be seen nearby the halide ion. The s-adenosylhomocysteine products are shown as sticks in colors matching their corresponding structure. N- and C-termini are labelled as black and white spheres, respectively. Split sites are labelled as spheres for *Bm*-MHT (light blue) and M.HhaI (light purple). (**B**) Topology diagrams from the Rossmann-fold core regions of *At*-MHT (blue) and M.HhaI (purple) methyltransferases. Beta sheets are shown as triangles, alpha helices are shown as circles, and loop regions are shown as lines. Split sites are shown as red crosses. The SAM and substrate binding regions are shown as black rectangles. For simplicity only secondary structure elements from the Rossmann-fold regions are included.

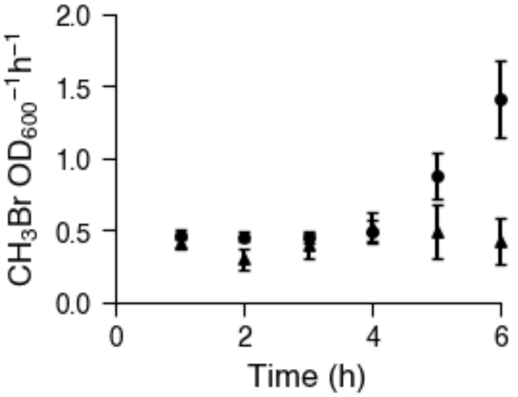


**Figure S2. $CH_3Br$ production from sMHT-113 at different temperatures.** The rate of $CH_3Br$ accumulation (μM/cell/hour) measured at different temperatures shows that the rate observed after 6 hours is significantly higher at 30 °C (squares) compared with 37 °C (triangles) ($p = 2.3 \times 10^{-3}$). Data represent the average of 3 biological replicates, with error bars representing +/- 1 standard deviation. Significance was evaluated using a two-sided, independent t-test.

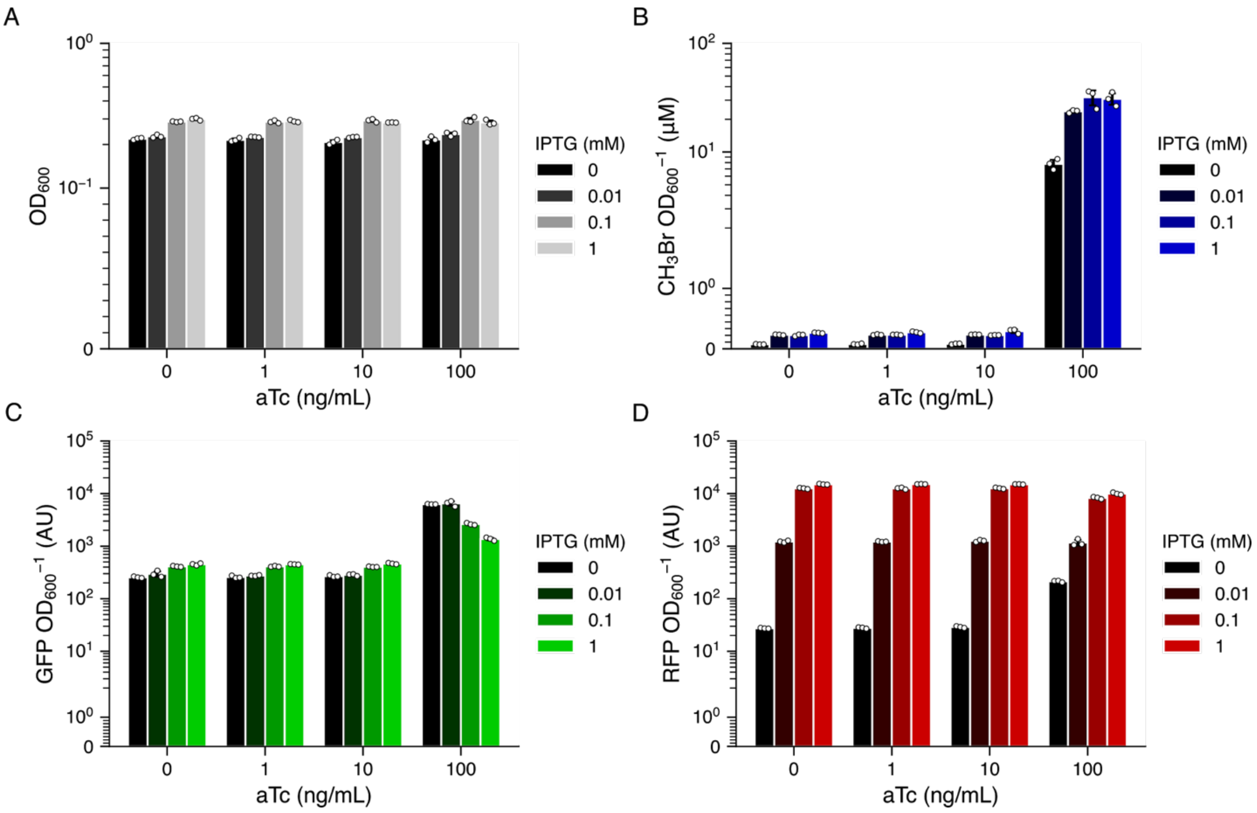


**Figure S3. Two-input AND-gate measurements. (A)** $OD_{600}$ values for cultures grown in the presence of different aTc (0, 1, 10, 100 ng/mL) and IPTG (0, 0.01, 0.1, and 1 mM) concentrations. All cultures grew to a similar $OD_{600}$ ($p > 0.01$). **(B)** $CH_3Br$ accumulation in the same cultures after 24 h. The relative $CH_3Br$ production was calculated as the ratio of $CH_3Br$ concentration to $OD_{600}$ to account for variability in cell growth. **(C)** GFP fluorescence in the same cultures after 24 h. GFP fluorescence was calculated as the ratio of green emission to $OD_{600}$ to account for variability in cell growth. **(D)** RFP fluorescence in the same cultures after 24 h. RFP fluorescence was calculated as the ratio of red emission to $OD_{600}$ to account for variability in cell growth.

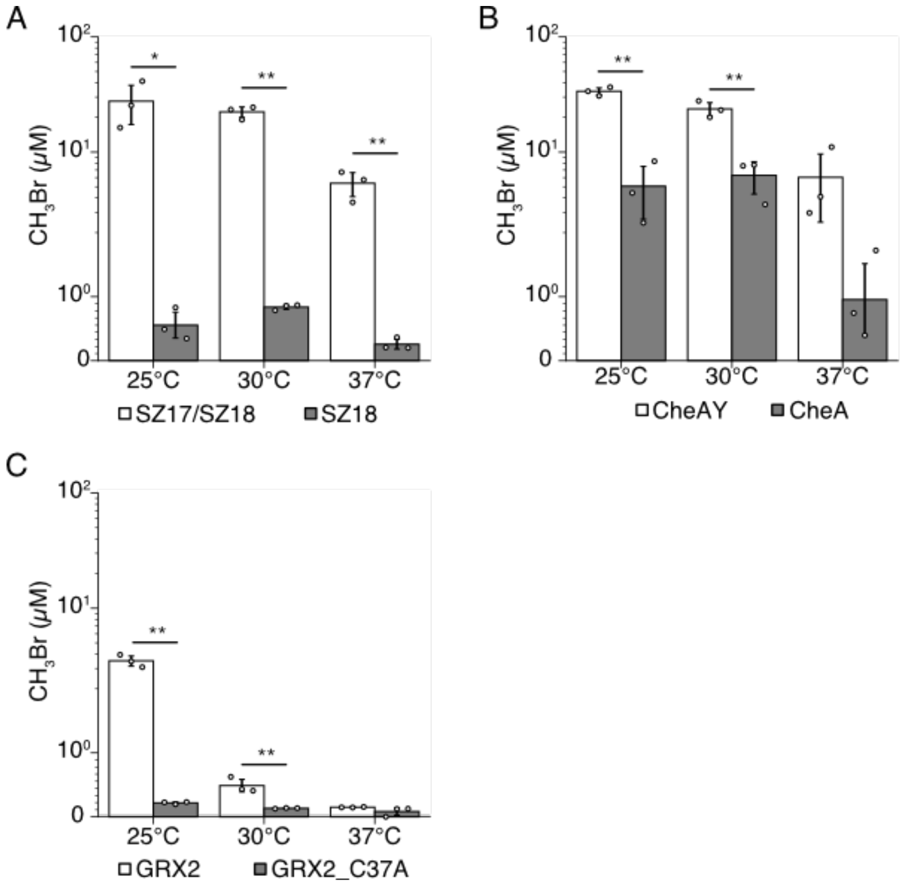


**Figure S4. $CH_3Br$ production by sMHT-113 fusions at varying temperatures.** $CH_3Br$ production at 25, 30, and 37°C by (**A**) sMHT-SYNZIP (SZ17/18) fusions, (**B**) sMHT-CheAY fusions, and (**C**) sMHT-GRX2 fusions. White bars represent constructs with both sMHT fragments tethered to interacting partner proteins. Gray bars represent constructs where F1 is expressed without fusion to the interacting protein partner (SZ18 and CheA in panels A and B) or where the protein partner's ability to dimerize has been removed via mutation (GRX2_C37A, panel C). All constructs showed a significant ($p < 0.05$) decrease in $CH_3Br$ production between dimerizing and non-dimerizing constructs at 25 and 30°C. Data represent the average of 3 biological replicates, with error bars representing +/- standard deviation. Significance values (*: $p < 0.05$, **: $p < 0.01$) were calculated using a two-sided, independent t-test.

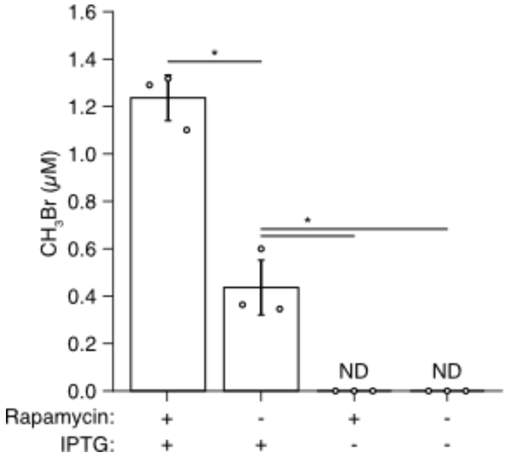


**Figure S5. $CH_3Br$ production by sMHT-FKBP/FRB fusions.** The effect of rapamycin (100 μM) and IPTG (100 μM) on $CH_3Br$ synthesis by cells containing a vector for LacI-controlled coexpression of F1-FKBP and FRB-F2. $CH_3Br$ production was measured following incubation at 37°C for 24 hours in closed vials. The gas production in the presence of both IPTG and rapamycin was significantly higher than that observed with IPTG or rapamycin alone ($p < 0.01$). Data represent the average of 3 biological replicates, with error bars representing +/- standard deviation. Significance values (***:** $p < 0.01$) were calculated using a two-sided, independent t-test.

### Supplementary Methods

**Gas production rates by sMHT-113-SYNZIP at varying temperatures**. To characterize $CH_3Br$ synthesis rates, *E. coli* MG1655 harboring a construct that constitutively expresses both sMHT-113-SYNZIP fragments, was grown to exponential phase in M63 media with 100 μg/mL streptomycin. 1 mL culture was diluted to an $OD_{600}$ = 0.1 in a capped vial, cells were incubated at the indicated temperatures for one hour, and indicator gas was measured using GC-MS. Following each measurement, vials were uncapped, the $OD_{600}$ was measured, and the vial was aerated (20 times with 50 mL of air each time) to purge $CH_3Br$ molecules in the gas and liquid phase. Each vial was recapped, incubated for an hour, and the sampling/purging was repeated.

**Gas production rates by sMHT-113 fusions at varying temperatures.** To evaluate the effect of temperature on $CH_3Br$ production by sMHT fused to SYNZIP, CheAY, and GRX2 partners, gas production by *E. coli* XL1 harboring each sMHT-partner fusion as well as the corresponding non-dimerizing control was measured. Cultures in M63 in sealed vials were prepared as in other gas production experiments (see *Methods*), protein expression was induced with IPTG (1 mM), and cultures incubated at 25, 30, or 37°C for 18 hours before measuring $CH_3Br$ accumulation.

**Gas production by sMHT-FKBP/FRB fusions +/- IPTG and rapamycin.** To assess the background production of $CH_3Br$ by sMHT-FKBP/FRB, cultures were prepared as previously described (see *Methods*). Briefly, overnight *E. coli* CS50 cultures expressing sMHT-113 FKBP/FRB fusions were diluted 1:100 into M63 media and grown to mid-log phase. IPTG (1 mM) and/or rapamycin (100 μM) was added, and cultures incubated for 24 hours at 37°C before $CH_3Br$ measurement by GC-MS.
